# Modulation of P2X4 pore closure by magnesium, potassium, and ATP

**DOI:** 10.1101/2021.05.16.444323

**Authors:** Kalyan Immadisetty, Josh Alenciks, Peter Kekenes-Huskey

## Abstract

The P2X4 receptor plays a prominent role in cellular responses to extracellular ATP. Through classical all-atom molecular dynamics (MD) simulations totaling 24 *µ*s we have investigated how metal-complexed ATP stabilizes the channel’s open state and prevents its closing. We have identified two metal-binding sites, magnesium (Mg^2+^) and potassium (K^+^), one at the intersection of the three subunits in the ectodomain (MBS1) and the second one near the ATP binding site (MBS2), similar to those characterized in Gulf coast P2X. Our data indicate that when Mg^2+^ and K^+^ ions are complexed with ATP, the channel is locked into an open state. Interestingly, irrespective of the number of bound ATP molecules, Mg^2+^ ions bound to the MBS2 resisted collapsing of the open state protein to a closed state by stabilizing the ATP-protein interactions. However, when Mg^2+^ in the MBS2 was replaced with K^+^ ions, as might be expected when in equilibrium with an extracellular solution, the interactions between the subunits were weakened and we found evidence of pore collapse. This collapse was apparent when fewer than two ATP were bound to MBS2 in the presence of K^+^. Therefore, the different capacities of common cations to stabilize the channel may underlie a mechanism governing P2X4 channel gating in physiological systems. This study provides structural insights into the differential modulation of ATP activation of P2X4 by Mg^2+^ and K^+^.

## 1 INTRODUCTION

Purinergic (P2X) receptors are ligand-gated cation channels that localize to the plasma and intracellular vesicle membranes (1). P2X4 receptors are expressed in almost all mammalian tissues (2). P2X4 receptors are implicated in regulating cardiac function, ATP-mediated cell death, synaptic strengthening, activation of the inflammasome in response to injury, multiple sclerosis (3–7), and neuropathic pain mediated by microglia (8–11). P2X4 is activated by adenosine triphosphate (ATP) (3, 12–14), during which the channel opens and allows the rapid flow of ions such as calcium (Ca^2+^), Mg^2+^, sodium (Na^+^), and K^+^ (14) between cellular compartments. Although the P2X4 receptor exhibits low Ca^2+^ permeability, it is often associated with triggering Ca^2+^-sensitive intracellular processes (14–16).

P2X4 is a homotrimer (17, 18), with each subunit consisting of a transmembrane domain (TMD) and an extracellular ectodomain (Fig. S1). Each TMD contains two TM helices (TM1 and TM2). The TM2 of each subunit together forms the TM pore that controls the gating of the channel (17, 18). P2X receptors are capable of binding up to three ATP molecules (19). Each ATP binds in an orthosteric binding site (Fig. S1) between the two neighboring subunits and facilitates a conformational transition from a closed (inactive) to an open (active) state (17, 18). Although mechanisms of P2X4 function are increasingly understood, several structural (role of intracellular fragments and pore dilation) and functional aspects (activation mechanism, allosteric modulation by agonists such as ivermectin, the number of ATP needed to activate the channel, desensitization, and the role of metal cations in modulating the ATP-mediated activation of the channel) of P2X4 channels are still elusive (20, 21). Knowledge about these processes is essential to understand how P2X4 function drives cellular responses. A pressing limitation is the lack of human resolution P2X4 structures to provide structural insights into its mechanisms. However, zebrafish P2X4 (zfP2X4) have been crystallized in several functionally important conformations (17, 22). zfP2X4 shares ∼59% sequence identity with human P2X4 (20) and has been shown to form functional homomeric channels with properties comparable to mammalian orthologs (23). Hence, zfP2X4 structures are reasonable surrogates for the modeling of human variants of this channel (20, 23).

Ions play a critical role in the normal functioning of many proteins (24–28); it is, therefore, no surprise that divalent cations including zinc, magnesium, copper, cadmium, silver, and mercury were reported to modulate P2X receptors. For instance, it was demonstrated that zinc and copper modulate rat P2X4 receptors differently, i.e., zinc potentiates, whereas copper inhibits ATP current (29, 30). Furthermore, it was demonstrated via site-directed mutagenesis the role of residue C132 in zinc potentiation (31), and residues D138 and H140 in copper inhibition in rat P2X4 receptors (31, 32). Similarly, it was reported that cadmium facilitates, whereas mercury inhibits ATP mediated currents in rat P2X4 receptors (33). Intriguingly, it was hypothesized that there are at least three metal-binding sites in P2X channels (33). Kasuya *et al*. reported an X-ray structure of Gulf Coast P2X receptor complexed with ATP and zinc (34), for which identified two metal-binding sites: MBS1 and MBS2. The MBS1 is located at the intersection of three isomers in the ectodomain, whereas the MBS2 is located near the ATP binding site between the two isomers. Zinc and magnesium bind to MBS1 and MBS2, respectively, and potentiates ATP current. Li *et al*. identified magnesium binding to MBS2 near the ATP binding domain in human P2X3 that strengthened ATP-protein interactions, which delay ATP unbinding and channel recovery from desensitization (35). Li *et al*. also proposed two different binding modes for magnesium, each in the presence and absence of ATP (35). It has been reported that cations binding to MBS2 in different P2X receptors have diversified roles (34–36). In Gulf Coast P2X, it was demonstrated that zinc bound to MBS1 potentiates ATP mediated currents, but not when bound to MBS2 (34). Riedel *et al*. reported that K^+^ and Na^+^ modulate human P2X7 receptor-operated single channel currents (37). The same study also reported a binding site for Na^+^on the extracellular side.

Despite the known dependencies of P2X4 function on metal cation binding, the structural mechanisms by which common ions like Mg^2+^ and K^+^ impact the channel function are less understood. All-atom molecular dynamics (MD) simulations have been used to investigate aspects of P2X4 function, and generally encompass short nanosecond level unbiased MD simulations or enhanced sampling MD simulations to probe P2X4 activation (38–45). To our knowledge, MD studies reported thus far have not examined how cations modulate ATP activation of P2X4, which is important for understanding channel function in mixed electrolyte solutions typical of cellular environments. Hence, we investigated the metal-binding sites, as well as the differential role of metal cations such as Mg^2+^ and K^+^, in modulating the ATP mediated activation of P2X4 via microsecond-length MD simulations (totaling 24 *µs*). We have identified two metal-binding sites MBS1 and MBS2, similar to gulf coast P2X (34), for which Mg^2+^ and K^+^ binding to the latter have different capacities to facilitate P2X4 pore closing.

## 2 RESULTS AND DISCUSSION

### 2.1 Mg^2+^ ions modulating ATP activation of P2X4

Three inter-subunit metal and ATP binding sites were identified in Gulf Coast P2X (34) and human P2X3 (35), which have been coined as the metal binding site 2 (MBS2) sites. To understand the role of Mg^2+^ binding at the MBS2 site in P2X4, we simulated five P2X4 open state structures. Three of the states have one equivalent of Mg^2+^ bound for each ATP: 3 ATP-3 Mg^2+^ (three ATP and three Mg^2+^ bound), 2 ATP-2 Mg^2+^ (two ATP and two Mg^2+^ bound), and 1 ATP-1 Mg^2+^ (one ATP and one Mg^2+^ bound). We also considered two cases with excess bound Mg^2+^: 1 ATP-3 Mg^2+^ (one ATP and three Mg^2+^ bound) and 2 ATP-3 Mg^2+^ (two ATP and three Mg^2+^ bound). (Fig. 1, Fig. S2). The latter were modeled in order to determine whether ATP was necessary to facilitate Mg^2+^ binding between the interfaced channel subunits. Three replicas of each system were simulated for 1 *µ*s each. All five systems stabilized after 400 ns as reflected in the protein backbone root mean squared deviations (RMSD) (Fig. S3A) that approached less than 4 Å in all cases (Fig. S3). We first investigated the stability of Mg^2+^ ions in the MBS2 sites. Our data suggest that Mg^2+^ were not stable in the absence of ATP in the neighboring nucleotide binding site. Specifically, for the 1 ATP-3 Mg^2+^ and 2 ATP-3 Mg^2+^ systems, two and one Mg^2+^ that were bound to the vacant MBS2 sites left the sites spontaneously (Fig. S4).

**Figure 1:**
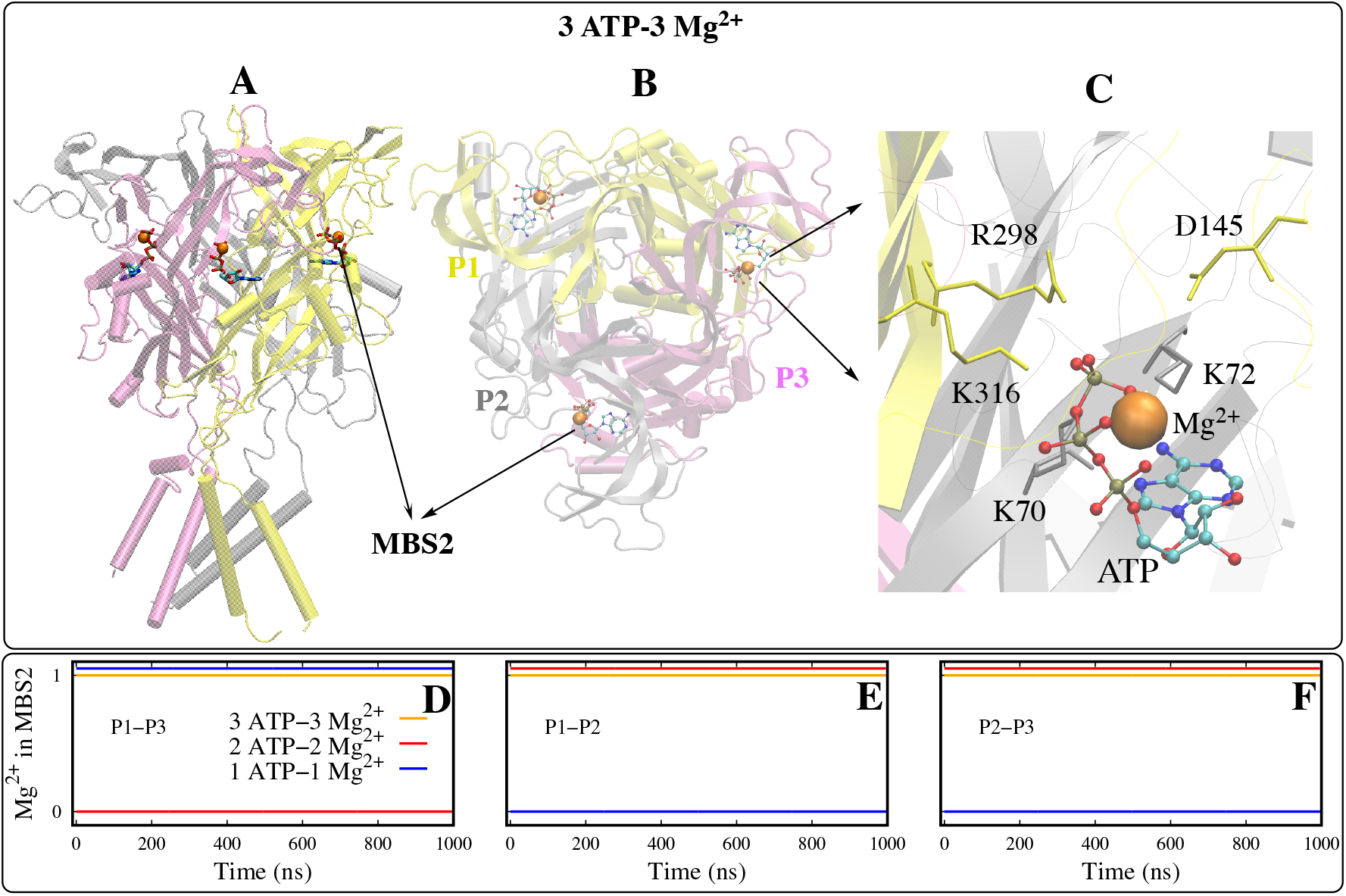
The open state P2X4 structure bound to three ATP and three Mg^2+^ (3 ATP-3 Mg^2+^). Mg^2+^ were docked into MBS2, which is located near the ATP binding site. P1, P2, and P3 isomers are colored yellow, grey, and magenta, respectively. Shown in panels A and B are the side and top views of the channel. The close-up of MBS2 is shown in panel C. Residues within 6 Å of Mg^2+^ are shown. Residues are colored to match the respective isomer. Mg^2+^ are shown as orange spheres and ATP is represented as a ball and stick model. (D–F) Time series of presence of Mg^2+^ in the the three MBS2 sites. 0 and 1 on Y-axis represent the absence and presence of Mg^2+^, respectively, in the MBS2 sites.

Activation of the channel by ATP opens the TM pore; pore opening is frequently measured by the positions of the L351 residue on each subunit, which borders the narrowest part of the TM pore (Fig. 2A) (19). Therefore, we report the radius of gyration (R_*g*_) of the residues L351 from each subunit (Fig. 2A) to assess the extent of the pore opening. The R_*g*_ values for all four systems (except for 1 ATP-3 Mg^2+^) were distributed between 4.5 and 6 Å, with an average of ≈5 Å that is comparable to the open state zfP2X4 crystal structure (≈5.3 Å). We did not observe significant differences in the distribution of R_*g*_ values between the systems simulated (Fig. 2B and Fig. S5). Hence, these data indicate that irrespective of the number of ATP and Mg^2+^ binding, the channel has not collapsed back to the closed state.

**Figure 2:**
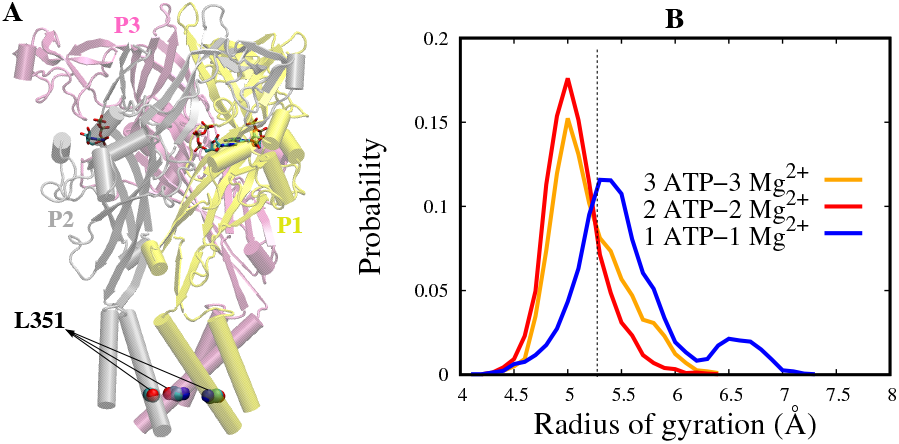
Probability density of the radius of gyration (R_*g*_). L351 of the three isomers were used for the calculation of R_*g*_ (A). Only C_*α*_ atoms of L351 were considered. Probability densities of Mg^2+^ bound systems are shown in panel B. The dotted vertical line represents the R_*g*_ of the open state zfP2X4 crystal structure.

### 2.2 K^+^ ions modulating ATP activation of the P2X4

The data in Section 2.1 suggest that the channel is stabilized in an open state irrespective of the number of bound ATP. Therefore, we speculated Mg^2+^ may exchange with monovalent ions like Na^+^ and K^+^, given that the concentration of Mg^2+^ is roughly 100-fold lower than the 154 mM intracellular concentration of K^+^ and extracellular concentration of Na^+^, respectively. To investigate this hypothesis, we simulated three P2X4 open state receptors, each docked with one, two, and three ATP molecules, respectively (Fig. S9) in the absence of Mg^2+^. During the simulation, solvated K^+^ ions infiltrated the MBS2 region and interacted with the ATP (Fig. 3A,B), as at least one K^+^ ion was bound in each MBS2 (Fig. 3D–F). Importantly, we found that MBS2 sites lacking ATP did not accommodate K^+^ ion, which was consistent with our findings using Mg^2+^.

**Figure 3:**
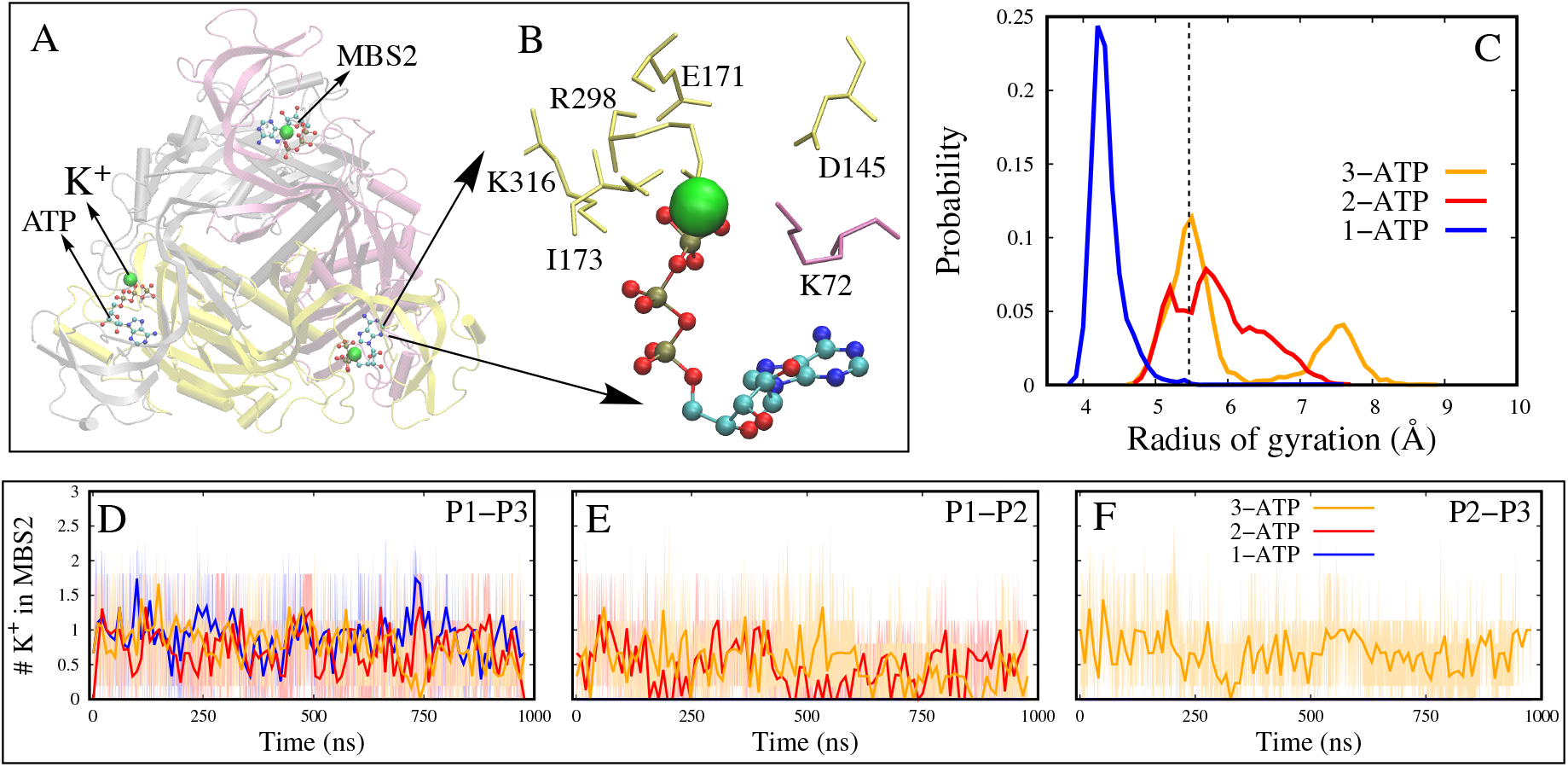
K^+^ binding in MBS2 site in P2X4. (A) K^+^ ions within 5 Å of ATP and protein in MBS2 are shown. (B) MBS2 between P1 and P3 is shown in close-up. Green spheres are K^+^ ions. ATP is shown as a ball and stick model. Residues within 5 Å of ions are shown. Residues are colored to match the respective isomers. (C) Probability density plots of the radius of gyration (R_*g*_) values of residue L351. The dotted vertical line represents the R_*g*_ of the open state zfP2X4 crystal structure. (D–F) The average number of K^+^ ions binding to MBS2s between the isomers P1,P3 (F), P1,P2 (G), and P2,P3 (H). Ions within 5 Å of both ATP and protein are estimated. 1-, 2-, and 3-ATP systems are colored blue, red, and orange, respectively. Average and standard of the three MD trials are shown.

To assess the impact of the number of ATP molecules and the binding of K^+^ ions on the opening/closing of the channel, we measured the pore radius by estimating the R_*g*_ of the TM pore residue L351 as explained in Sect. 2.1. While the 2- & 3-ATP systems resisted the pore collapse as was observed for the Mg^2+^-bound systems, the TM pore of the 1-ATP system collapsed within the first 5 ns of these simulations. The average pore radius for the 1-ATP system was ≈4.2 Å (Fig. 2C), whereas for the 2- and 3-ATP systems it was ≈5.5 Å (Fig. 2C), except for one 3-ATP trail (Fig. 2C). These data suggest that the substitution of K^+^ for Mg^2+^ weakens the coupling between channel subunits, such that the channel collapses when fewer than two ATP are bound. We speculate that the 1-ATP system conformation obtained in our simulation may resemble a desensitized state in which ATP is still bound but the TM pore is closed; full deactivation would requite ATP to unbind completely (35, 46).

### 2.3 Mechanisms of K^+^ vs. Mg^2+^ channel closing

To investigate the potential mechanisms coupling the MBS2 sites to channel opening, we performed dynamic cross-correlation analysis (DCCA) to identify the correlations between dynamic regions of the receptor. Correlation coefficient matrices were computed for each system to identify regions of strong positive (red) or negative (cyan) correlations (Fig. S7). In general, the matrices were similar among the Mg^2+^ bound systems, which suggests that channel dynamics are comparable irrespective of the number of bound Mg^2+^-ATP equivalents. However, to better highlight the similarities, we provide a correlation of each subunit residue with respect to several functionally important residues: K70, D145, and L351 (Fig. 4). Residue K70 contributes to the MBS2 within a given subunit, whereas D145 lines the MBS2 site of an adjacent subunit. In Fig. 4A we show that K70 of one subunit is anticorrelated with the D145 of the same subinit (Fig. 4B, D), indicating that these two residues are pulled in opposite directions. We additionally observed that the TM pore residue L351 is negatively correlated with the remainder of the subunit (e.g., the MBS2 residues) (Fig. 4B, D).

**Figure 4:**
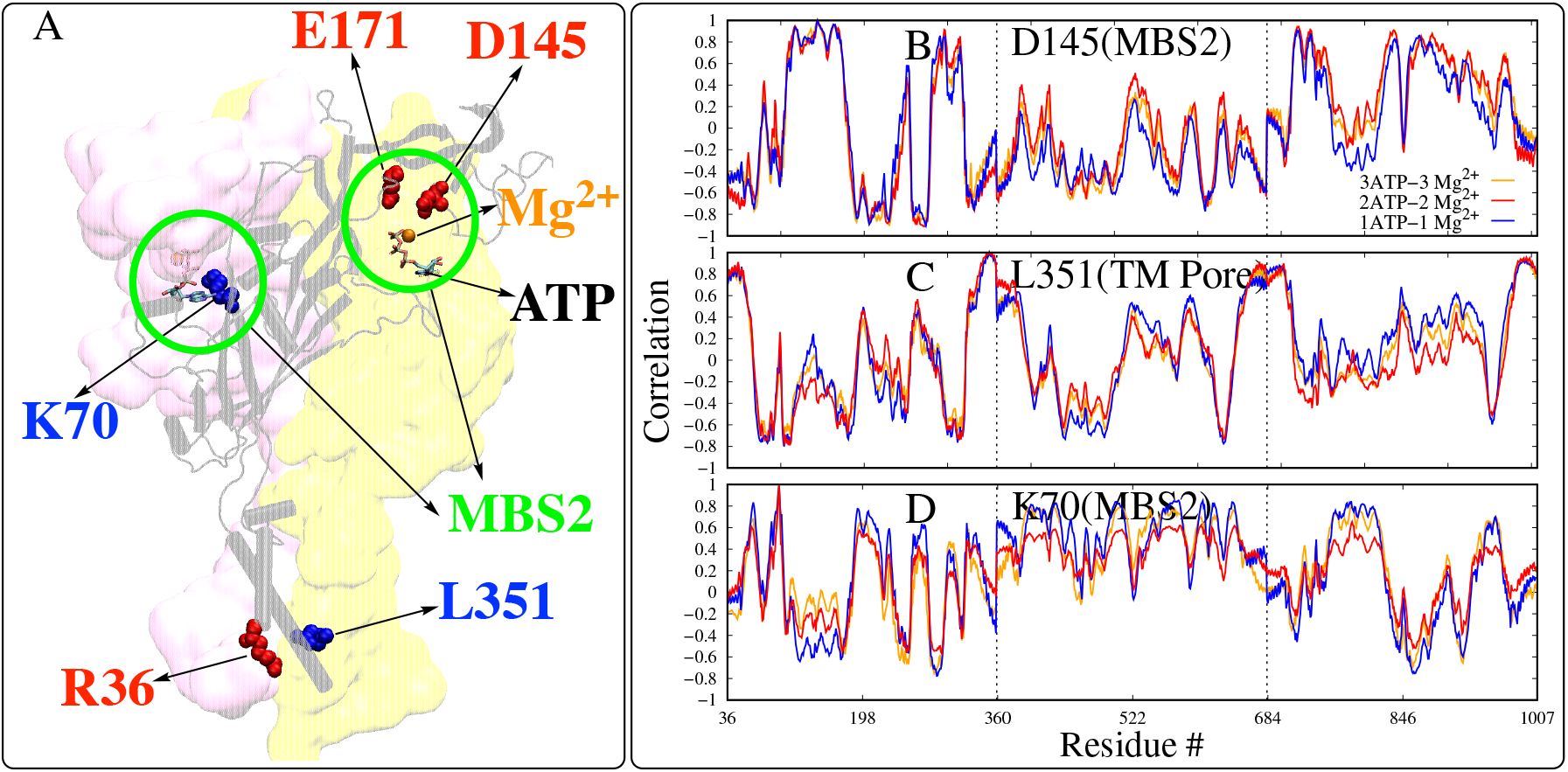
Inter-residue correlations in Mg^2+^ bound systems. (A) Residues of interest that belong to the isomer (colored grey and represented as a cartoon) are shown. The blue-colored residues are positively correlated with each other and the red and blue residues are anticorrelated. ATP is shown as sticks and Mg^2+^ bound in the MBS2 site is shown as orange spheres. The two MBS2 sites are circled in green. (B–C) Correlation plots of a select few residues of interest are shown on right. The dotted lines separate the three subunits. -1, 0, and 1 on the y-axis refers to anticorrelation, no correlation, and positive correlation, respectively.

Given our observation that the channel partially closes when one equivalent of ATP is bound in the absence of Mg^2+^, we hypothesized that replacing Mg^2+^ with K^+^ in MBS2 would weaken the dynamic coupling between the MBS2 and the pore regions. Indeed, the DCCA data for the 2- and 3-ATP/K^+^ bound systems (Fig. S7 and Fig. S12) were similar to the Mg^2+^ bound systems (Fig. 4 and Fig. S7). In contrast, the DCCA data were significantly different in the presence of a single bound ATP compared to the 3- and 2-ATP systems, respectively (Fig. S7 and Fig. S12). As an example, the correlation of the MBS2 site residue D145 with the L351 of a subunit is ≈7% in the 3 ATP-3 K^+^ system, whereas their motions were strongly negatively correlated (≈ −50%) between the same residues in the same subunit in the 1 ATP-1 K^+^ system (Fig. S12). Overall, our K^+^ bound simulations suggest that the ATP-K^+^ bound complex weakly stabilizes the subunit interfaces, such that losing two K^+^ is sufficient for the system to collapse back to a closed state.

### 2.4 Mg^2+^ vs. K^+^ coordination of ATP-protein interactions

To determine the basis of the different correlation patterns reported in Section 2.3, we examined how the protein coordinates the ATP/metal complexes at the MBS2 sites. It is evident in all ATP-bound cases that the adenine base of ATP directly interacts with one subunit (i.e., the subunit colored yellow in Fig. 5A, B), whereas its phosphate backbone interacts with residues D145 and E171 of a neighboring subunit (i.e., the subunit colored grey in Fig. 5A, B) via metal cations. In other words, the bound ATP bridges two interfaced subunits. Interestingly, waters appear to mediate the interactions between metals and residues D145 and E171. Analogous sites are implicated in coordinating the binding of metal ions in the MBS2 site in Gulf coast P2X (34) and hP2X3 (35). To quantify this coordination, we assessed the distribution of oxygens contributed by the protein and associated waters about Mg^2+^ and K^+^ cations bound at the MBS2 site (Fig. 5). The radius of the first oxygen shell near Mg^2+^ is at ≈2 Å and comprises four water oxygens, with two oxygens from the ATP phosphate and minor (< 1) contributions from the receptor. Visual inspection of the binding site indicates the bound waters bridge Mg^2+^ with residues D145 and E171. We report similar oxygen distributions for K^+^ as well, albeit the radius of the first oxygen shell was considerably larger (≈3 Å). Although both ions seem to utilize similar binding partners, bound Mg^2+^ may afford tighter coordination at the MBS2 site, which could serve to effectively lock adjacent channel subunits into an open configuration. K^+^ ions, in contrast, more weakly interface the subunits, which potentially permit channel closure. Hence, the bound waters might contribute to a water-mediated allosteric network stabilizing the P2X4 channel open state, similar to water-mediated hydrogen-bond networks shown to govern the conformational dynamics of other proteins such as aurora kinase A (47).

**Figure 5:**
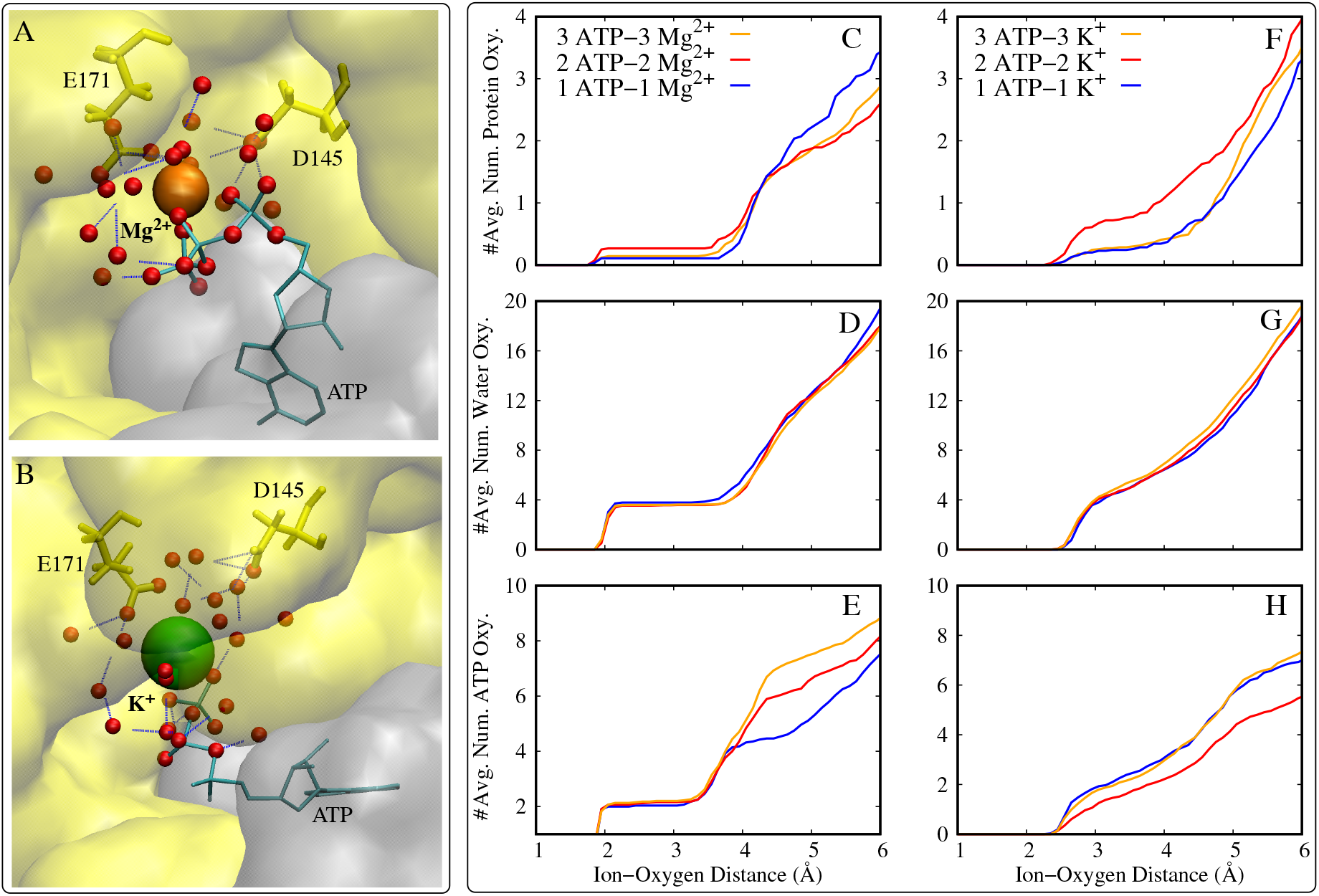
Metal cations coordinating the ATP-protein interactions in the MBS2 sites. (A) MBS2 3 ATP-3 Mg^2+^ system; (B) MBS2 in 3 ATP-K^+^ bound system. The two neighboring subunits are shown in yellow and grey and in surface representation. Oxygen atoms (belonging to water, protein, and ATP) within 6 Å of the ions are shown as red spheres. Hydrogen bonds are shown as blue broken lines. Mg^2+^ and K^+^ are shown as orange and green spheres, respectively. ATP is shown as cyan and in stick representation. Residues D145 and E171 are shown as yellow sticks. ATP connects the subunit colored grey to the one in yellow (i.e., to residues D145 and E171) via metal ions and water molecules. (C–H) Oxygen coordination around Mg^2+^ (C–E) and K^+^ (F–H) ions binding in the MBS2. Integrated amino acid, water, and ATP oxygen distribution are shown separately for each ion. The average number of oxygens around a single Mg^2+^ or K^+^ ion is shown. All three MD trials of each system were combined for these calculations.

## 3 CONCLUSION

In this report, we studied the modulation of ATP activation of P2X4 by metal cations via all-atom MD simulations. We identified two metal-binding sites similar to other P2X receptors, such as Gulf coast P2X. The first metal-binding site (MBS1) was found at the intersection of three isomers in the upper part of the vestibule and the second metal-binding site (MBS2) resided near the ATP binding site. We investigated how the binding of Mg^2+^ versus K^+^ at MBS2 modulates ATP mediated channel closing. Our data suggest Mg^2+^ ions stabilize the coupling between channel subunits and thereby prevent the collapse of the open state to a closed state, irrespective of the number of ATP molecules binding to the channel. It has been reported that the ATP-Mg^2+^ complex delays the human P2X3 receptor’s recovery from desensitization (35), hence bound Mg^2+^ in P2X4 may contribute to a long recovery from desensitization. In contrast, the exchange of Mg^2+^ with K^+^, which is expected when more highly concentrated K^+^ is in thermodynamic equilibrium with low concentration Mg^2+^, permits channel closing when a single equivalent of ATP is bound. We therefore speculate that weakening of the subunit coupling, e.g., via Mg^2+^ replacement with K^+^ facilitates the collapse of the channel. Interestingly, it was found for a related purinergic receptor, P2X7, that one ATP was sufficient to activate the channel even in the absence of Mg^2+^ (48), which suggests that purinoreceptors may utilize different mechanisms for ion-dependent regulation. Our simulations also suggest that MBS1 readily binds metal cations (K^+^) to bridge the interactions between its three subunits, but does not appear to be sensitive to the metal identity at MBS2. For this reason, they may contribute to stabilization of the trimer interface. Altogether, our study provides insight into metal-mediated interactions in the P2X4 channel that may contribute to its channel gating properties in multi-electrolyte cellular environments.

## 4 METHODS

### 4.1 Molecular dynamics simulations

The open state crystal structure of zfP2X4 bound with three ATP molecules (PDB: 4DW1 (22)) was utilized for this study. Initially, the open state zfP2X4 crystal structure was obtained from the protein data bank (49) and all waters were removed. Two and one ATP bound open state zfP2X4 systems were generated by removing ATP from the three ATP bound crystal structure. All three systems were further processed and built for the MD simulation using the CHARMM-GUI web server (50, 51). Namely, each system was placed in the 1-palmitoyl-2-oleoyl-sn-glycero-3-phosphocholine (POPC) bilayer and solvated using TIP3P water (52). K^+^ and chloride (Cl^−^) ions were added to balance the charges as well as to attain a physiological concentration of 0.15 M. Each system contained one P2X4 protein, ≈42,425 TIP3P water molecules, ≈388 POPC lipid molecules, 0.15 M KCl, and 3/2/1 ATP molecules. The size of each system was ≈155 × 160 × 170 Å^3^ and the total number of atoms in each system is ≈195,116. The amber force field (ff12SB) (53) was used to treat the entire system. All three systems described above were simulated (three replicas for each system) with Mg^2+^ ions docked near the site of ATP binding as described in the references (34, 35). For the 2-ATP and 1-ATP systems, Mg^2+^ was only docked to the sites that contain ATP. Additionally, 2-ATP and 1-ATP systems were generated but with Mg^2+^ docked in all three domains irrespective of the presence of ATP molecules. Overall, five Mg^2+^ docked open state zfP2X4 systems were modeled for MD simulations; a 3-ATP system with Mg^2+^ found at each docking site, two 2-ATP systems, and two 1-ATP systems. The 2-ATP and 1-ATP had one system each with Mg^2+^ docked in all three domains and one system each where Mg^2+^ only docked in the domains where ATP is present. Mg^2+^ was removed from the input structures in order to model the K^+^-bound configurations; however, the K^+^ ions were not docked into the MBS2 sites but rather spontaneously coordinated from the surrounding water box during the MD simulations.

Each system was energy minimized for 100,000 steps using a conjugate gradient algorithm (54), and further relaxed in a multistep procedure (which spans for ≈1.5 ns), wherein the lipid tails, protein side chains, and backbone were restrained and then released in a step-wise manner as explained elsewhere (50) in the NVT ensemble. Furthermore, production simulations were conducted under periodic boundary conditions in NPT ensemble. Three replicas of each system were simulated, each replica for ∼ 1 *µs* (1*µs* × 3 trials × 3 systems = 9*µs*). AMBER16 (55) was used for conducting the MD simulations. A 2 fs time step was used for the initial relaxation as well as for the follow-up production simulations. The temperature was maintained at 310 K using a Langevin thermostat and the Nose-Hoover Langevin piston method was used to maintain a 1 atm pressure (56, 57). The nonbonded interactions were cut-off at 12 Å and the particle mesh Ewald (PME) method (58) was used to treat the long-range electrostatics. Trajectories were saved every 20 ps. The SHAKE algorithm was used to constrain the hydrogen bonds (59).

All simulations were conducted on a local GPU cluster and using XSEDE resources. Data analysis was conducted using VMD plugins (60) and the cpptraj module of Amber (61). Five data points per each ns were used for analysis. Simulation input files and generated data are available upon request. Dynamic cross-correlation maps were generated to understand the correlated motions of the protein. Cpptraj (61) package of Amber was used for users to conduct this analysis and only C_*α*_ atoms of the protein were used to calculate the correlation coefficients. All three MD trials of each system were combined and a correlation matrix was generated for the entire protein.

## 5 AUTHOR CONTRIBUTIONS

KI and JA conducted the simulations, analyzed the data, and wrote the manuscript. PKH designed the project, analyzed the data, and wrote the manuscript.

## 6 ACKNOWLEDGMENTS

Research reported in this publication was supported by the Maximizing Investigators’ Research Award (MIRA) (R35) from the National Institute of General Medical Sciences (NIGMS) of the National Institutes of Health (NIH) under grant number R35GM124977. This work used the Extreme Science and Engineering Discovery Environment (XSEDE) (62), which is supported by the National Science Foundation grant number ACI-1548562.

## 7 SUPPLEMENTARY MATERIAL

**Figure S1:**
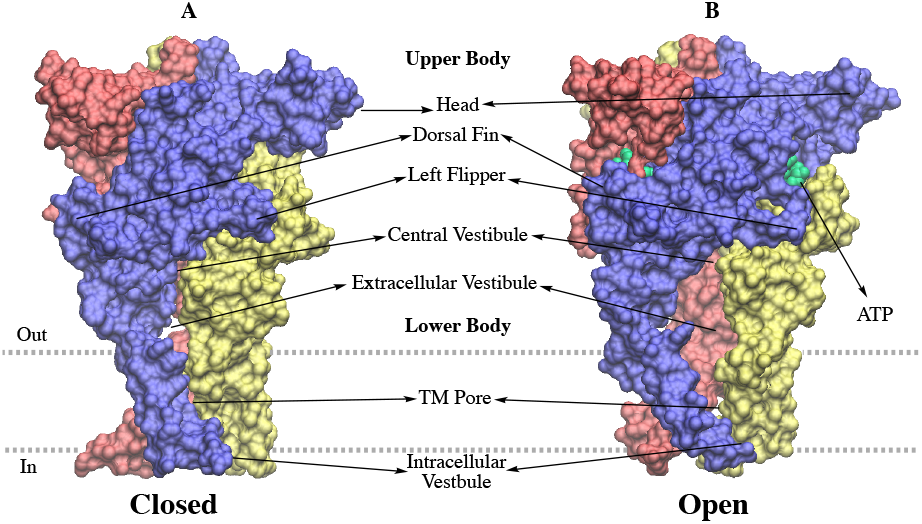
Closed (A) and open state (B) crystal structures of zfP2X4. The three isomers are colored blue, red, and yellow, respectively. The closed structure is in apo form and ATP is binding in the open state structure between the isomers. ATP is shown in green. The TM pore, which is formed by the TM domain of each isomer, is the narrowest part of the channel. The TM pore is closed when deactivated and open when activated.

**Figure S2:**
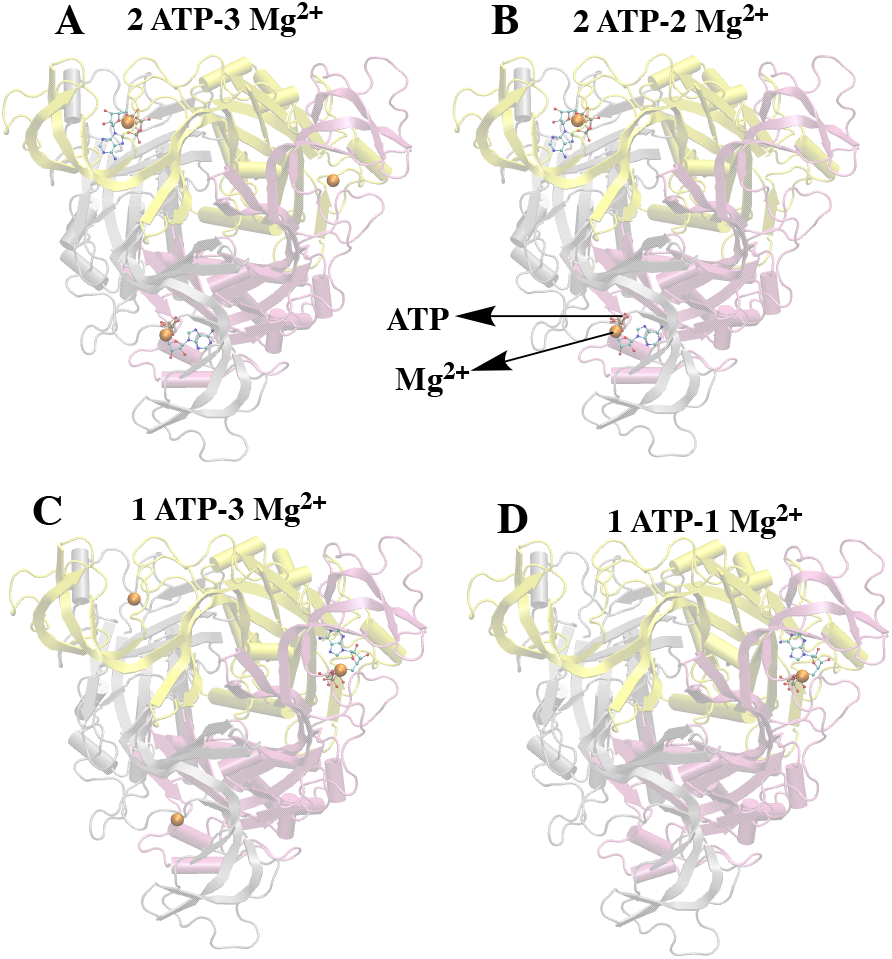
The remaining four p2×4 structures were studied with Mg^2+^ docked in MBS2. Top views of all four structures are shown. (A) 2 ATP-3 Mg^2+^ structure has two ATP and three Mg^2+^ ions, (B) 2 ATP-2 Mg^2+^ structure has two ATP and two Mg^2+^ ions, (C) 1 ATP-3 Mg^2+^ structure has one ATP and three Mg^2+^ ions, and (D) 1 ATP-1 Mg^2+^ structure has one ATP and one Mg^2+^ ions. Mg^2+^ is shown as orange spheres and ATP is represented as a ball-and-stick model.

### 7.1 MBS1 binding

The number of K^+^ binding in MBS1 was estimated. On average, two to three K^+^ ions were consistently present in MBS1 in the 3-ATP and 2-ATP systems, whereas in the 1-ATP system they were reduced from three to one in the course of the simulation (Fig. 3E). We hypothesize that this is due to the collapse of the channel in the 1-ATP system, thus collapsing MBS1. Studies have reported that zinc (Zn^2+^) binding to MBS1 potentiates ATP-mediated currents in the Gulf coast P2X channel through allostery (34). To some extent, we have observed K^+^ binding MBS1 playing a similar role as Zn^2+^. For instance, a greater number of K^+^ ions are binding in this site when two or more ATP are binding in the MBS2. This indicates that K^+^ and Zn^2+^ are working in correlation and that K^+^ help stabilizes the open state structure in the presence of ATP, which is needed to facilitate greater ion/current flow. Additionally, we have identified that K^+^ ions were occupying the MBS2 and interacting with the ATP (Fig. 3A). Furthermore, we estimated that at least one K^+^ ion was bound in the MBS2 (Fig. 3F–H). Additionally, we identified a correlation between ATP binding to the channel and K^+^ ions binding to the MBS2, i.e., when no ATP binds to an ATP binding site, no K^+^ ions were binding to the respective MBS2, and vice versa. For instance, in the 3-ATP system, three ATP molecules bind to the three ATP binding sites and subsequently, we observed K^+^ ions binding to all three MBS2s. However, with the 2-ATP system K^+^ ions were bound near the two ATP binding sites that were occupied by ATP but not near the ATP binding site that was not occupied by ATP (i.e., the site between domains P2 and P3) (Fig. 3 F–H). Similarly, in the 1-ATP system, K^+^ was binding in the MBS2 adjacent to the site occupied by ATP (i.e., between the domains P1 and P3) and absent in the other two MBS2 sites. This is similar to what we have observed in the Mg^2+^ bound simulations.

**Figure S3:**
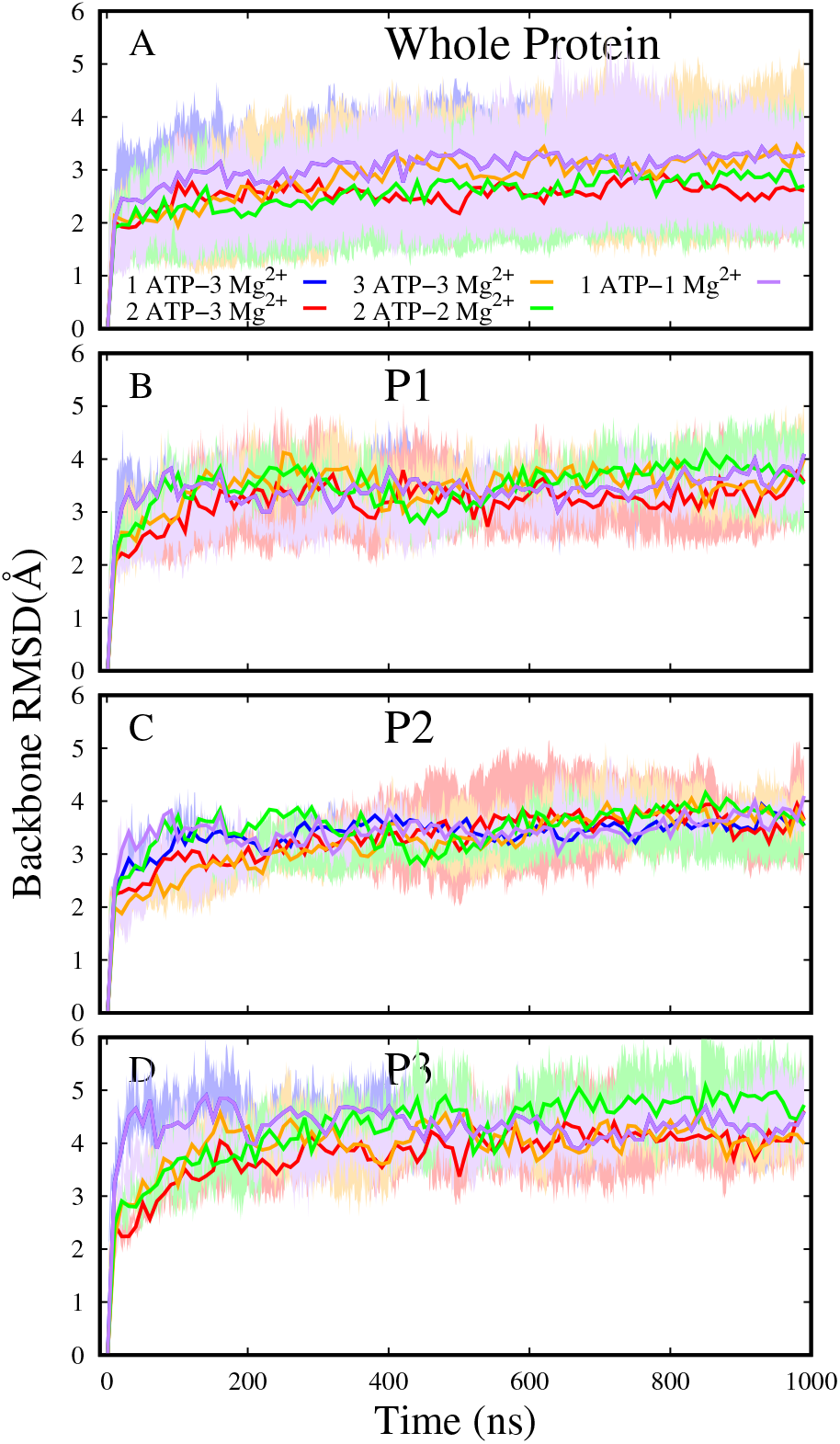
Backbone RMSD vs. Time for Mg^2+^ bound systems. RMSD of the whole protein is shown in A and of the three isomers in panels B–D. The average and standard deviation of all three MD are shown.

**Figure S4:**
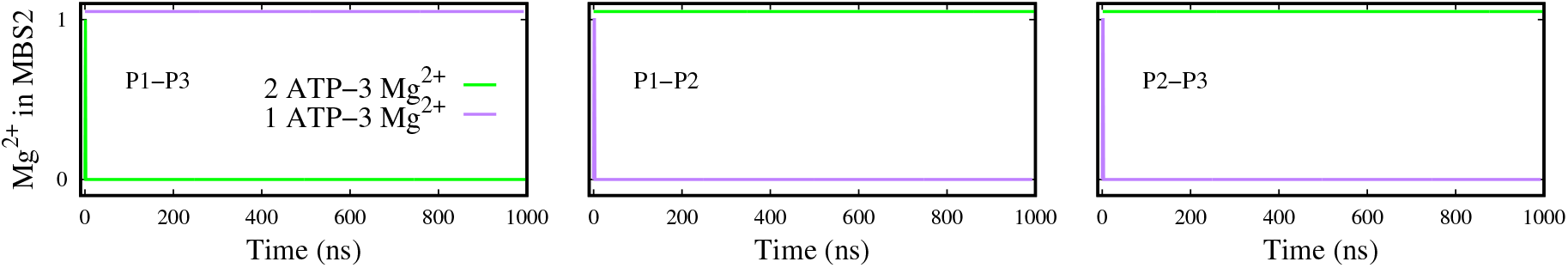
Mg^2+^ in MBS2 sites of 2 ATP-3 Mg^2+^ and 1 ATP-3 Mg^2+^ systems. The three columns represent Mg^2+^ in the three MBS2 sites in the channel. Mg^2+^ left the MBS2 sites in the absence of ATP in 1 ATP-3 Mg^2+^ and 2 ATP-3 Mg^2+^ systems.

**Figure S5:**
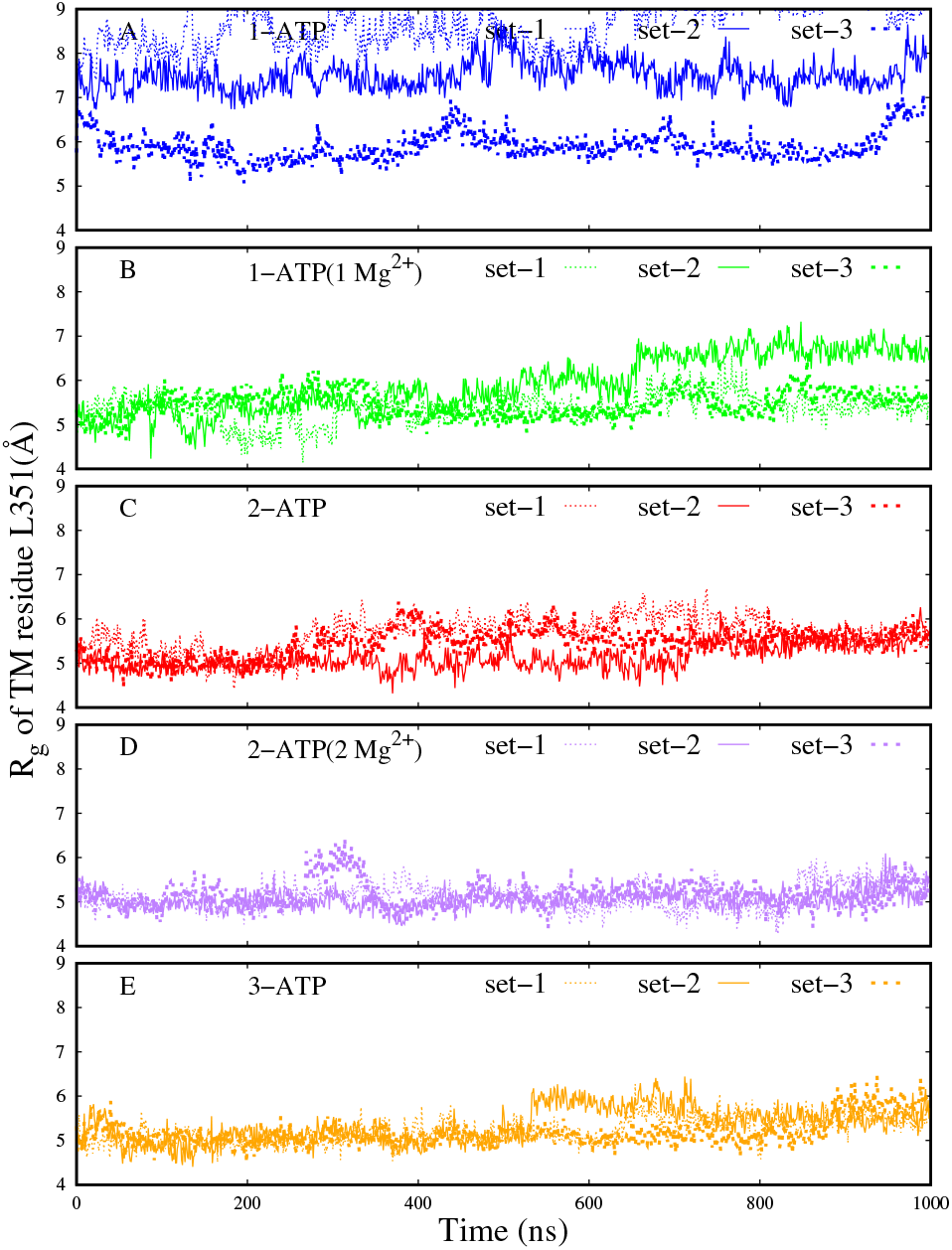
Radius of gyration vs. Time for Mg^2+^ bound systems. All three MD trials of each system are shown separately.

**Figure S6:**
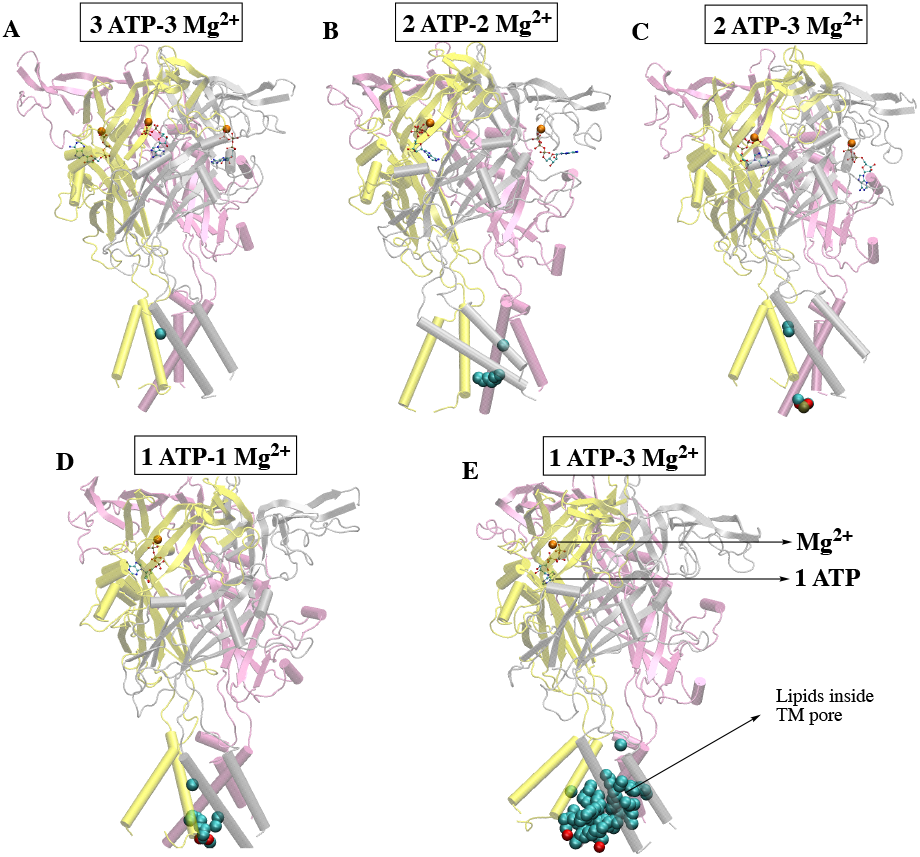
Representative MD snapshots of all five Mg^2+^ bound P2X4 systems. Lipids inside the TM pore are shown.

**Figure S7:**
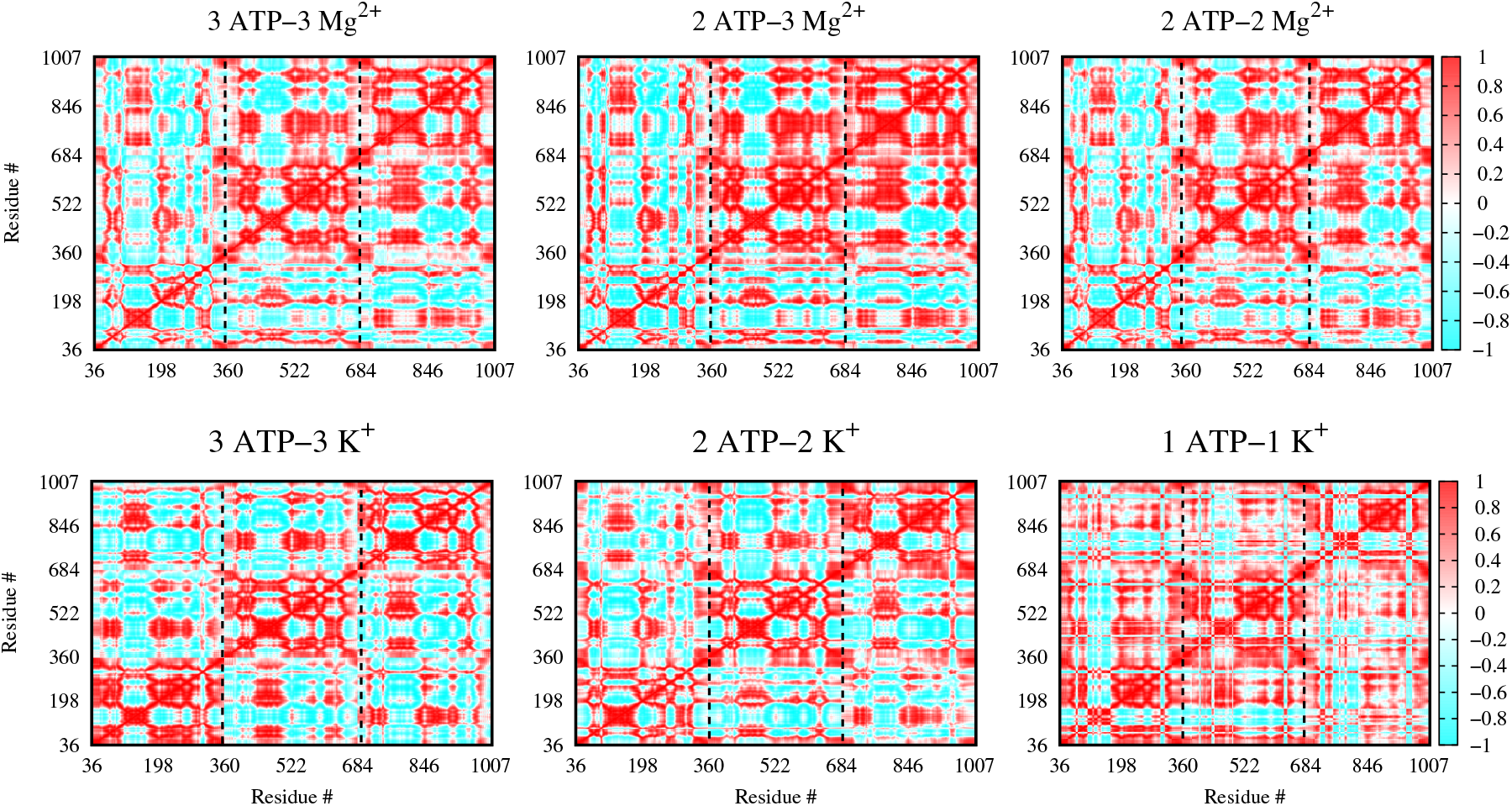
Dynamic cross-correlation heat maps of Mg^2+^ and K^+^ bound systems. All three MD trials of each system were combined for this analysis; only C_*α*_ atoms were considered. The color bar on the right represents the extent of cross-correlation; -1, 0, and 1 represent negative correlation (cyan), no correlation (white), and positive correlation (red), respectively. Cpptraj (61) of the amber package was used for this analysis.

**Figure S8:**
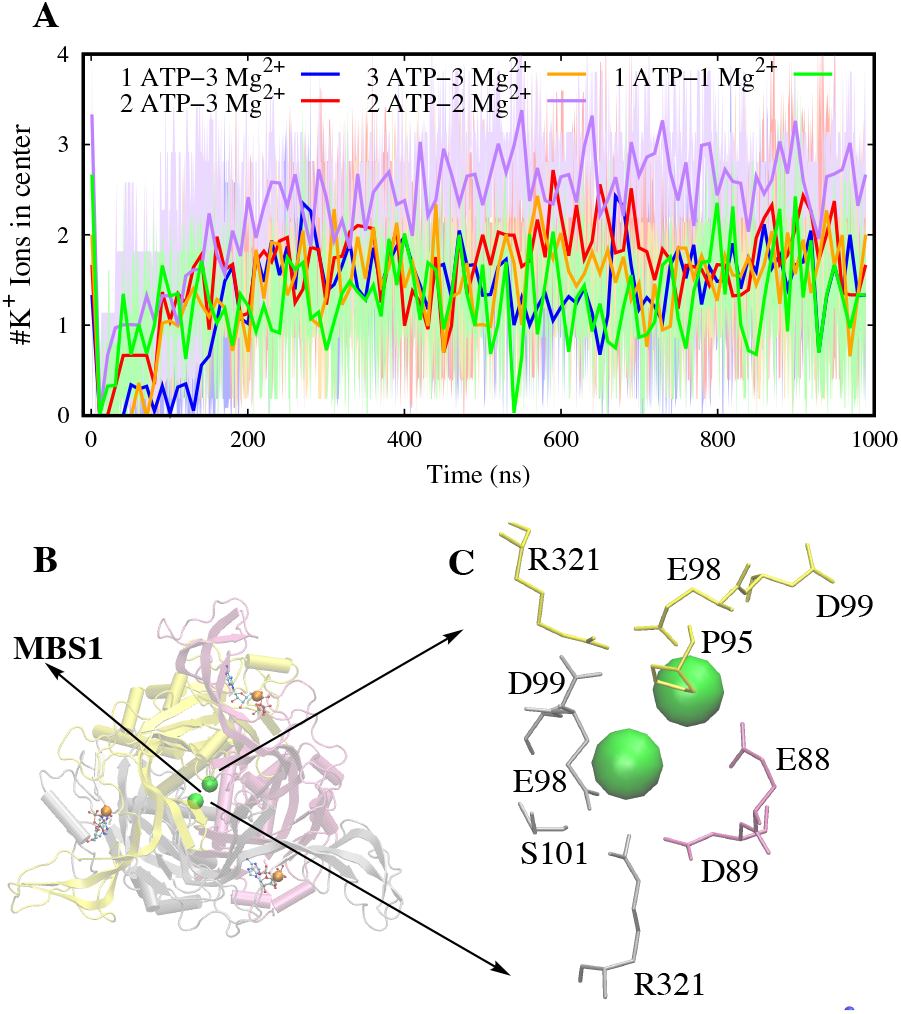
K^+^ ions in MBS1 in Mg^2+^ bound systems. The # of K^+^ found in the MBS1 site as a function of simulation time is shown. Average and standard deviation of three MD trials are shown. (B) MD snapshot of the 3 ATP-3 Mg^2+^ system with K^+^ binding to MBS1 is shown. (C) Close-up of MBS1 and residues within 5 Å of K^+^ are shown. Residues are colored according to the isomer they belong to. K^+^ ions are shown as green-colored spheres.

**Figure S9:**
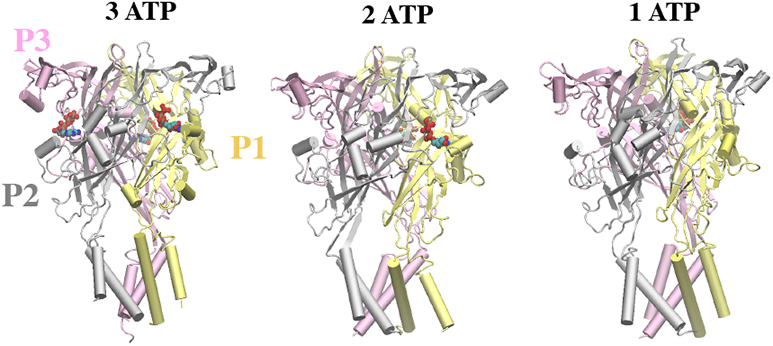
The three zfP2X4 structures studied. P1, P2, and P3 domains are colored yellow, grey, and magenta, respectively. ATP molecules are shown as ball-and-stick representations. 1-ATP: ATP binds between P1 and P3 domains; 2-ATP: ATP binds between P1, P2, and P1, P3 domains. Side views of all three structures are shown.

**Figure S10:**
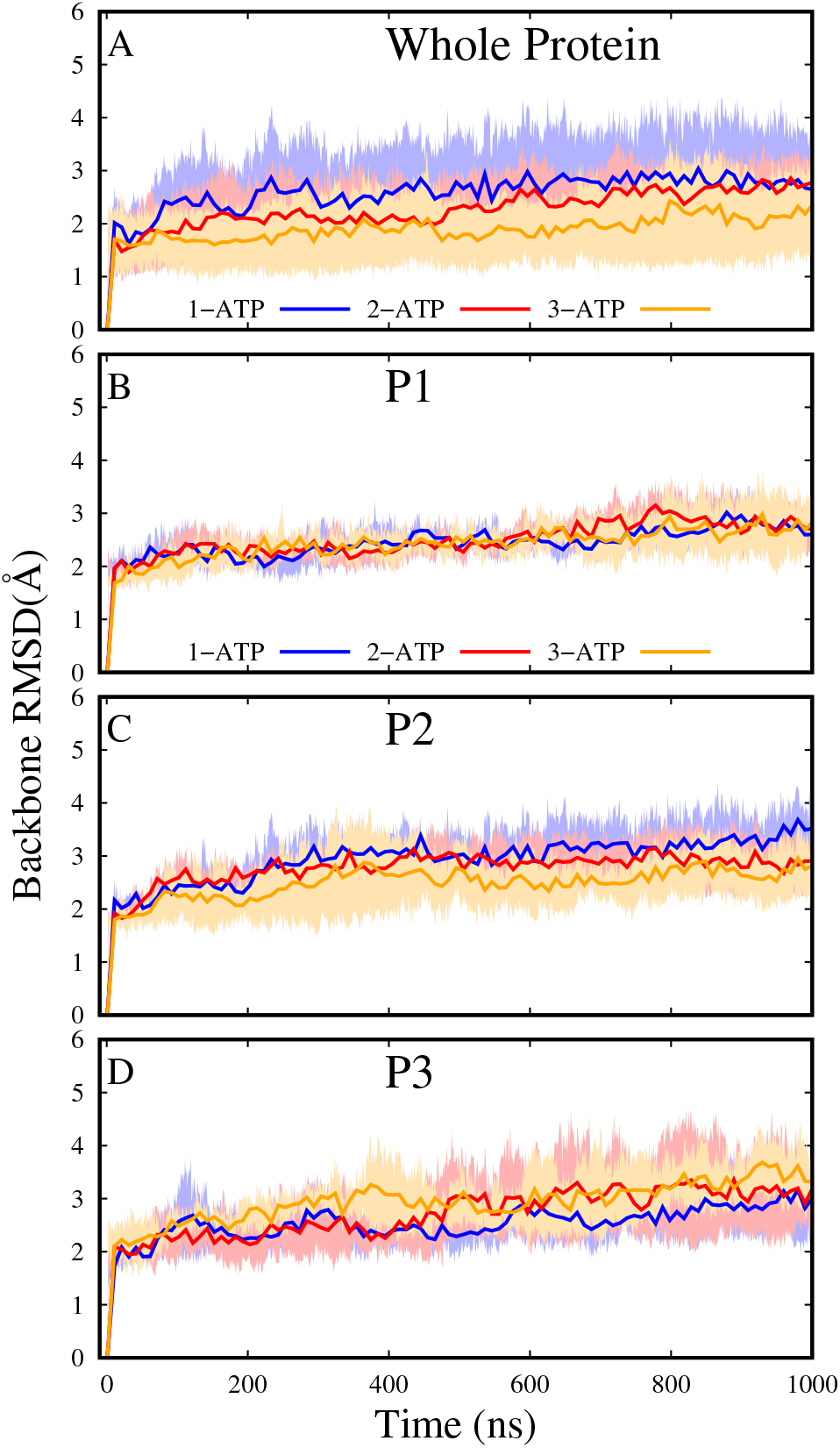
Backbone RMSD vs. Time. **A** RMSD of whole protein. **B-D** RMSDs of the three isomers. The average of three MD trials of each system is shown and the error bars (i.e., standard deviation) are shaded.

**Figure S11:**
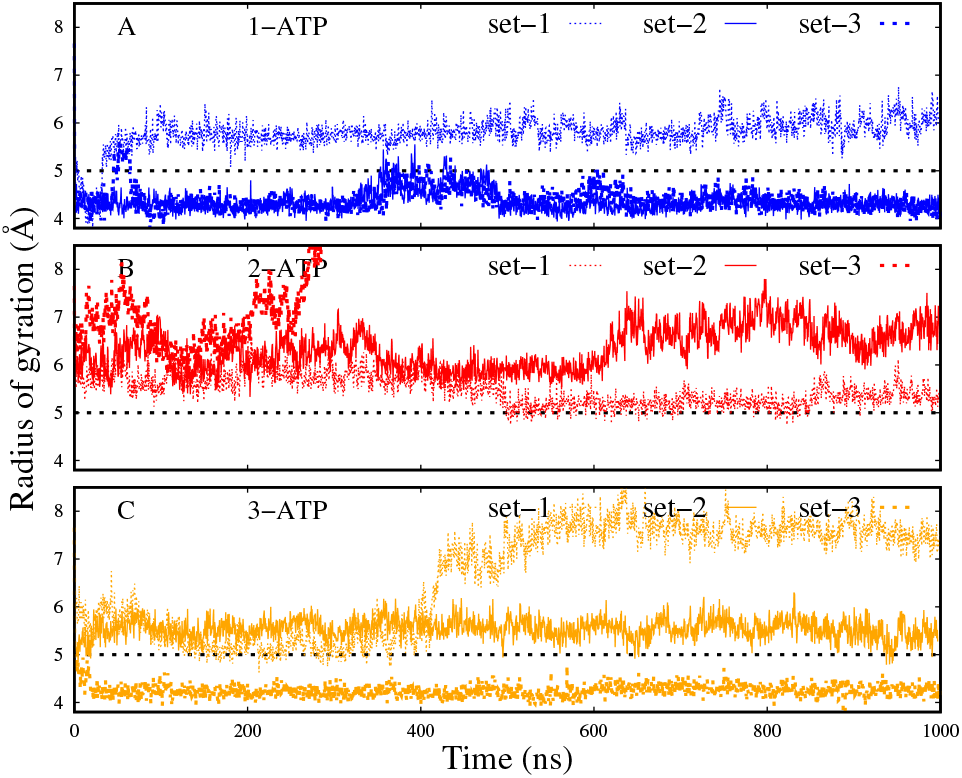
R_*g*_ vs. time of residue L351. 1-, 2-, and 3-ATP systems are colored blue, red, and orange, respectively. The dotted horizontal line at Y=5 separates 1-ATP system from the other two systems. R_*g*_ of 1-ATP systems were less than 5 Å, except for one outlier (set-1). On the other hand, the R_*g*_ of 2- & 3-ATP systems are greater than 5 Å, except for one outlier in the 3-ATP system (set-3).

**Figure S12:**
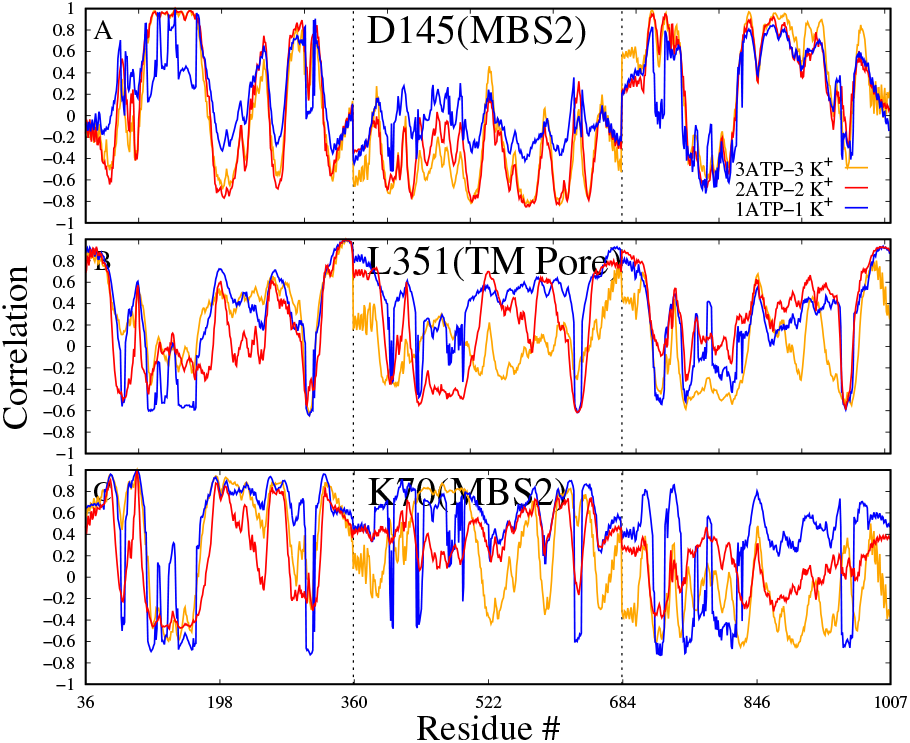
Correlation plots of select few residues of interest of K^+^ bound systems. The dotted lines separate the three monomers. -1, 0, and 1 on the y-axis refers to anti-correlation, no correlation, and positive correlation, respectively.

**Figure S13:**
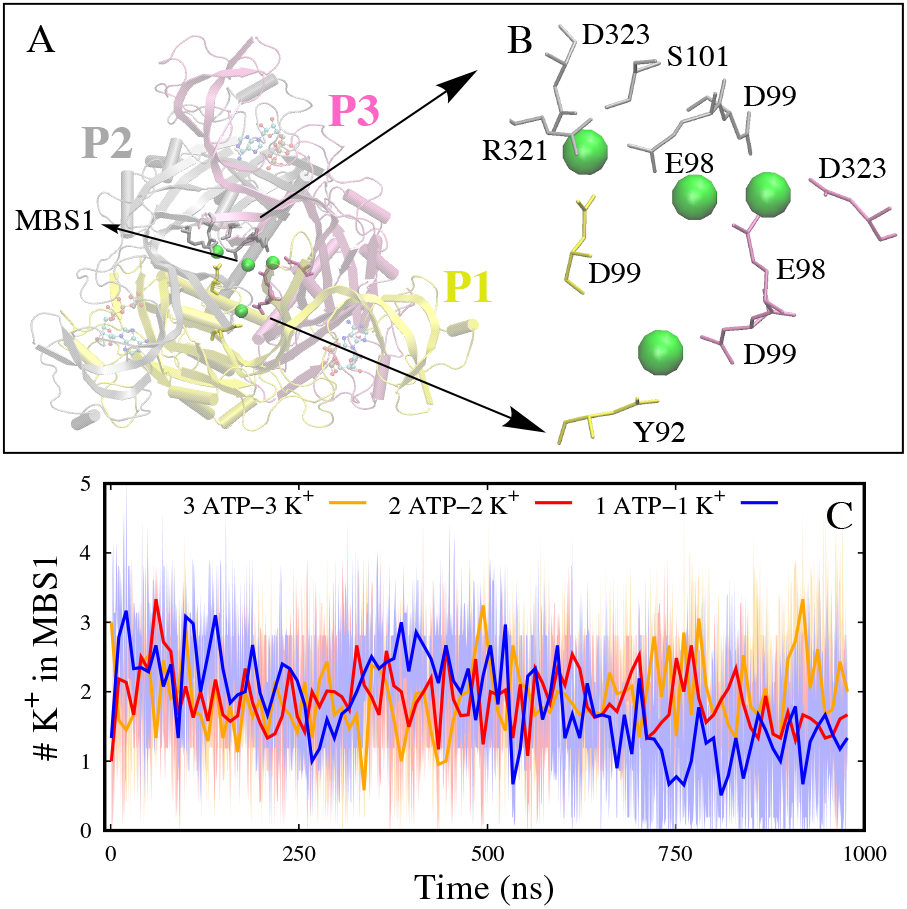
K^+^ ions binding in MBS1 coordinating the three isomers inside the vestibule. **A** Top view is shown. **B** MBS1 close-up view. **C** Average number of K^+^ ions binding in MBS1. Ions within 8 Å of all three isomers inside the vestibule are considered for these calculations.

We have observed K^+^ ions occupying the MBS1 in all five Mg^2+^ bound systems studied (Fig. S8) and coordinating the three isomers via interacting with the polar (such as S101 and Y99) and negatively charged (such as E98, D99, and D323) residues of the protein (Fig. S8B, C). Irrespective of the number of ATP binding to the channel, on average 2–3 K^+^ were binding to the MBS1 (Fig. S8A). This is a highly conserved site across the P2X4 receptors and residues that are interacting with the zfP2X4 are similar to other P2X receptors (34).

K^+^ ions were also identified in MBS1 in the three K^+^ bound systems simulated (Fig. S13).

## ACRONYMS

Ca^2+^: calcium. 2
Cl^−^: chloride. 7
K^+^: potassium. 1–7, 12, 15, 17, 18
MBS2: metal binding site 2. 3
MD: molecular dynamics. 1, 2, 4, 6, 7, 13–16
Mg^2+^: magnesium. 1–7, 12–15, 18
Na^+^: sodium. 2, 4
R_*g*_: radius of gyration. 3, 4, 16
RMSD: root mean squared deviations. 3, 13
TMD: transmembrane domain. 2
zfP2X4: zebrafish P2X4. 2–4, 7, 12, 15, 18
Zn^2+^: zinc. 12

